# Multiplex Embedding of Biological Networks Using Topological Similarity of Different Layers

**DOI:** 10.1101/2021.11.05.467392

**Authors:** Mustafa Coşkun, Mehmet Koyutürk

## Abstract

Network embedding techniques, which provide low dimensional representations of the nodes in a network, have been commonly applied to many machine learning problems in computational biology. In most of these applications, multiple networks (e.g., different types of interactions/associations or semantically identical networks that come from different sources) are available. Multiplex network embedding aims to derive strength from these data sources by integrating multiple networks with a common set of nodes. Existing approaches to this problem treat all layers of the multiplex network equally while performing integration, ignoring the differences in the topology and sparsity patterns of different networks. Here, we formulate an optimization problem that accounts for inner-network smoothness, intra-network smoothness, and topological similarity of networks to compute diffusion states for each network. To quantify the topological similarity of pairs of networks, we use Gromov-Wasserteins discrepancy. Finally, we integrate the resulting diffusion states and apply dimensionality reduction (singular value decomposition after log-transformation) to compute node embeddings. Our experimental results in the context of drug repositioning and drug-target prediction show that the embeddings computed by the resulting algorithm, Hattusha, consistently improve predictive accuracy over algorithms that do not take into account the topological similarity of different networks.

## 1 Introduction

Network models are commonly used to represent sets of biological interactions and associations in a network context[1]. The nodes of these networks represent biological entities, including genes, proteins, processes, functions, diseases, etc. The edges represent interactions and/or associations in different contexts, ranging from physical interactions to functional and statistical associations [14].

As more biological network data, along with advanced machine learning algorithms become available, these algorithms are increasingly applied to prediction tasks in biomedical applications [3]. These machine learning algorithms often require vector-space representations of features [16]. A common solution to this problem is in the form of node embeddings, which map the nodes of a network into a latent lower dimensional vector space [22]. Network embeddings are shown to be highly effective for various downstream tasks, including disease gene prioritization, prediction of drug response, and prediction of protein function [25].

In the context of biological networks, the interactions/associations between pairs of entities can represent a multitude of relationships. For example. the relationships between protein pairs include physical interactions, sequence similarity, regulatory interactions, functional similarity, shared pathway membership, etc [10]. Different networks with a common node set are often organized into multiplex networks, where each layer in the network represents a different type of relationship among the nodes [11]. This motivates the development of algorithms to compute node embeddings for multiplex biological networks [5].

Multiplex network embedding techniques aim to derive strength from a broad range of data sources by integrating multiple networks with a common set of nodes [5,18,21,26,27]. These algorithms, illustrated in Figure 1, usually compute diffusion states for the nodes of individual networks [9]. They then integrate the resulting diffusion states using probabilistic modeling [5], linear dimensionality reduction [5], or neural networks [27]. Algorithms that aim to generalize computation of diffusion states to multiplex networks have been also developed [23], but these algorithms treat all networks equally and do not take into account the differences between the topology or sparsity patterns of different networks.

**Fig 1:**
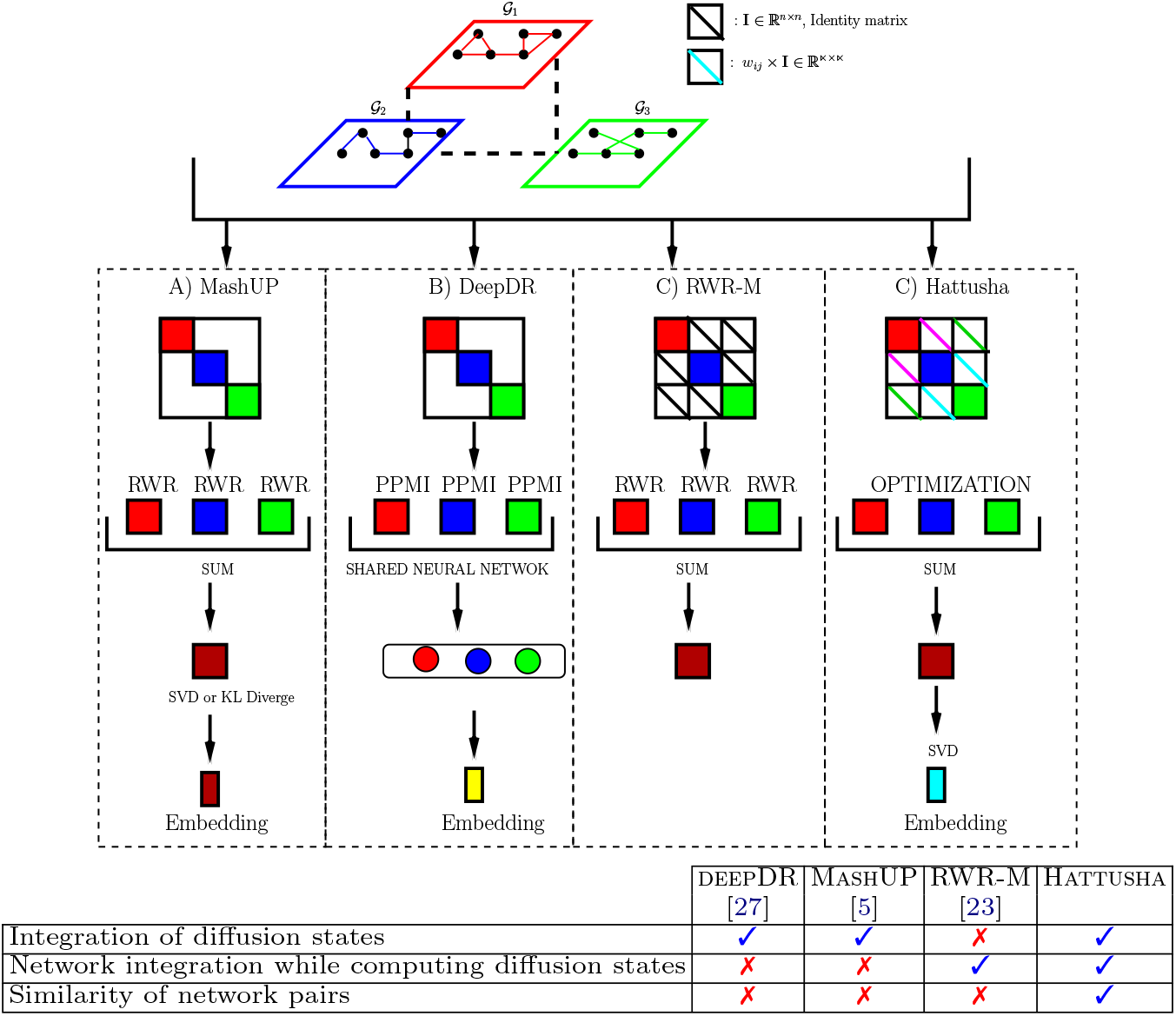
Existing approaches and the proposed approach to multiplex network embedding. MashUP and deepDR compute diffusion states independently on each network and then integrate diffusion states using dimensionarly reduction/learning algorithms, while RWR-M treats all networks equally while computing diffusion states on integrated networks. Hattusha integrates networks while computing diffusion states and considers similarity of network pairs while integrating networks.

We propose Hattusha, a multiplex network embedding algorithm that utilizes the topological similarity of pairs of layers in the multiplex network to compute diffusion states. Hattusha represents the computation of diffusion states as an optimization problem, with an objective function that is composed of intra-network smoothness, inter-network smoothness, and fidelity to the seed vector (diffusion states are computed by using each node as a seed vector). In this formulation, inter-network smoothness between pairs of layers is enforced based on the topological similarity of these networks. To quantify topological similarity between a pair of networks, we use Gromov-Wassertein Discrepancy (GWD) [6], a measure that is adopted from the literature on optimal transport [19]. To solve the resulting optimization problem, we use Chebyshev polynomials to accelerate the iterative computation [7]. To assess Hattusha’s performance in the context of relevant biomedical applications, we use Drug Repositioning/Repurposing and Drug-Target Interaction Prediction as benchmark problems. We systematically compare the predictive performance of the embeddings produced by Hattusha against a deep learning based approach [27], a model-based approach [5], and a multiplex network diffusion approach that does not account for topological similarity [23]. Our results show that Hattusha consistently outperforms existing approaches on both problems.

## 2 Methods

### Notation and Problem Definition

A multiplex network with *k* layers is a collection of *k* networks with a common set of nodes, but different edge sets, i.e., 𝒢_1_ = (𝒱, ℰ_1_), 𝒢_2_ = (𝒱, ℰ_2_), …,𝒢_*k*_ = (𝒱, ℰ_*k*_). Here, 𝒱 represents the common set of *n* = |𝒱| nodes and ℰ_*i*_ represents the set of edges among the nodes of *i*-th network, where |ℰ_*i*_| = *m*_*i*_. We denote the adjacency matrix of the *i*th network as **A**_*i*_ ∈ ℝ^*n*×*n*^. We use the notation **P** to denote the column-normalized stochastic matrix derived from adjacency matrix **A**.

Multiplex network embedding is the problem of computing a vector-space representation (embedding) for the nodes in a low-dimensional space, such that the proximity of the embeddings in this space will capture the topological proximity/similarity of nodes across these multiple networks.

### Existing Algorithms for Computing Multiplex Node Embeddings

The current state-of-the-art in multiplex network embedding is illustrated in Figure 1. Cho et al. [5] develop MashUP, demonstrating that the following framework is effective for computing node embeddings in multiplex biological networks:

1. For each node *q* ∈ 𝒱, compute diffusion state 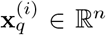 for all 1 ≤ *i* ≤ *k*, where 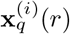 quantifies the proximity/similarity between *q* and *r* ∈ 𝒱 in 𝒢_*i*_.
2. Apply dimensionality reduction to these *k* × *n* diffusion states to compute node embeddings.

MashUP computes diffusion states individually on each network using standard random walk with restarts. Subsequently, it uses a multinomial logistic model along with a KL-divergence based loss function to compute low-dimensional representations that best approximate the profile vectors. An important drawback of MashUP is that it does not integrate the networks while computing diffusion states. A more recent algorithm, deepDR [27], uses a similar strategy in that it first computes individual embeddings and then integrates these embeddings across multiple networks using deep learning.

A recent algorithm that aims to directly compute diffusion states on multiplex networks is RWR-M [23]. RWR-M generalizes random walks to multiplex networks by replicating the *k* networks as multiple layers and adding inter-network edges between nodes that correspond to each other in different layers. RWR-M uses two hyper-parameters: (i) *δ* ∈ (0, 1), the damping factor of standard random walks that is used to tune the locality of the walk, and (ii) *λ* ∈ (0, 1), an additional parameter that is used to tune the frequency of passages between different networks. Given these two parameters, the adjacency matrix of the integrated multiplex network is defined as follows:

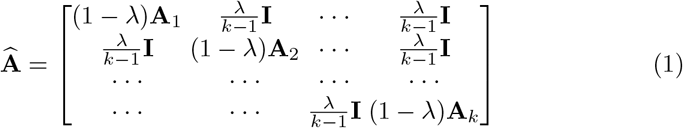

Here, **I** ∈ ℝ^*n*×*n*^ denotes the identity matrix, which is used to teleport from one layer to another through two corresponding nodes on different layers. Now for each node *q* ∈ 𝒱, multiplex random walk with restarts (RWR-M) is defined as:

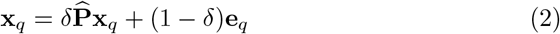

where **e**_*q*_ ∈ ℝ^*kn*^ denotes the seed vector with **e**_*q*_((*i* − 1)*n* + *q*) = 1*/k* for 1 ≤ *i* ≤ *k* and **e**_*q*_(.) = 0 for all other entries and 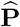 denotes the column-normalized stochastic matrix of **Â**. This linear system can be solved using the standard power method, i.e., by setting **x**_*q*_ = **e**_*q*_ and iteratively updating **y** until convergence [23]. The resulting vector **x**_*q*_ ∈ ℝ^*kn*^ is a concatenation of the desired network-specific diffusion states, i.e., 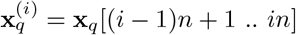 for 1 ≤ *k* ≤ *n*.

### 2.1 Proposed Approach: Weighted Transitions Between Layers

Existing approaches to multiplex network embedding have an important limitation: Although the variable density of the networks being integrated, as well as the relative similarity of different groups of networks, have a significant effect on the performance of downstream prediction tasks [12], all layers are treated equally. We address this issue by assigning weights to quantify the topological similarity of network pairs, and using these weights to control how different networks inform each other while computing diffusion states.

As described above, our objective is to first compute diffusion state vectors 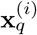 for all *q* ∈ 𝒱 and 1 ≤ *i* ≤ *k*. For a given *q* ∈ 𝒱, we set up *k* identical seed vectors 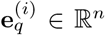 as 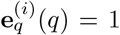 and 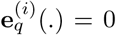 otherwise for 1 ≤ *i* ≤ *k*.

We then formulate the computation of these *k* diffusion state vectors 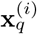 as an optimization problem:

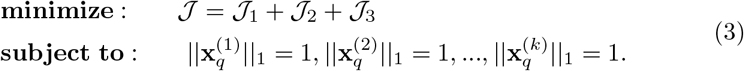

The objective function of this problem has three components: fidelity to the seed vector (𝒥_1_), intra-network smoothness (𝒥_2_), and inter-network smoothness (𝒥_3_).

The first two components of the objective function reflect the standard formulation for random walks [20], summed over all networks:

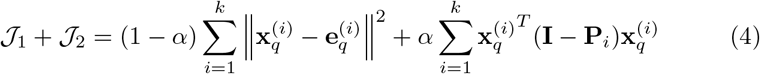

Here, **P**_*i*_ denotes the column-normalized stochastic matrix of network 𝒢_*i*_ and *α* ∈ (0, 1) is the damping factor. The third component of the objective function accounts for inter-network smoothness:

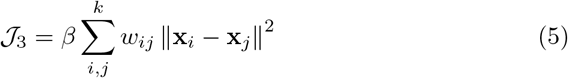

Here, *β* ∈ (0, 1) is a regularization parameter and *w*_*ij*_ ∈ (0, 1) for all 1 ≤ *i, j* ≤ *k* are weights that control the correspondence between pairs of networks. Observe that, in this formulation, pairs of networks that are assigned a larger weight are subject to more pressure to have similar diffusion states. In Section 2.2, we describe how these weights can be computed using Gromov-Wasserstein Discrepancy. Using these weights, we construct 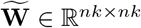 as:

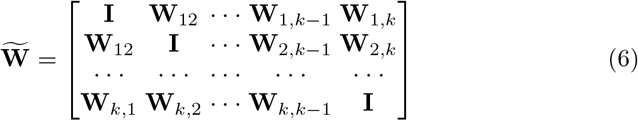

where **W**_*ij*_ = *w*_*ij*_ × **I**.

Now letting 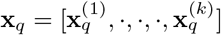 and 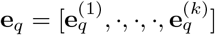 as in the previous section, we can rewrite the objective function as:

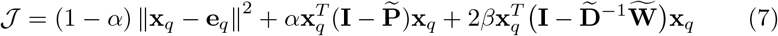

where 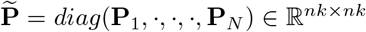 denotes the aggregated stochastic matrix and 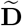 denotes the degree matrix of 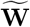.

By taking the derivative of 𝒥 with respect to **x**_*q*_, we obtain:

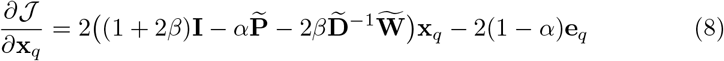

Since 𝒥 is a quadratic function of **x**_*q*_, we set the update rule as 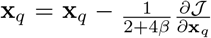 [17], and obtain:

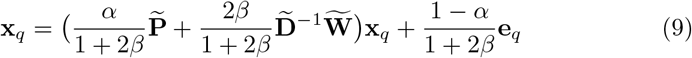

#### Lemma 1.

*The update rule for* RWR-M, 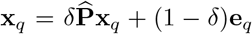, *is a special case of Equation (9)*.

*Proof*. By setting 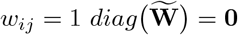 and 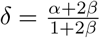, Equation (9) is reduced to Equation (2).

Note that the coefficient matrix of Eq. (9) exhibits numerical properties that allow implementation of Chebyshev acceleration to compute **x**_*q*_. For this reason, we implement Hattusha using Chebyshev acceleration [7]. The resulting algorithm and experimental results on the runtime improvement provided over standard power iteration are shown in the Supplementary Material.

### Computing Node Embeddings from Diffusion States

Once we compute **x**_*q*_ for each *q* ∈ 𝒱, we aggregate the diffusion states across networks to compute 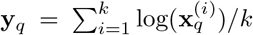. We then organize the aggregated profile vectors into an *n* × *n* profile matrix **Y** = [**y**_1_ **y**_2_ … **y**_*n*_] and apply Singular Value Decomposition (SVD) on **Y** to compute lower-dimensional node embeddings [5].

### 2.2 Computing Inter-Network Edge Weights via Graph Matching

We assess the similarity of pairs of networks using Gromov-Wasserstein Discrepancy (GWD), an optimization framework that is developed to compute the optimal transport between two sample spaces [19], and recently applied to the quantification of the topological (dis-)similarity between graphs [6].

Let **P**_*i*_ and **P**_*j*_ denote the column-normalized transition matrices for layers 𝒢_*i*_ and 𝒢_*j*_ of a multiplex network. The Gromov-Wasserstein Discrepancy (GWD) of 𝒢_*i*_ and 𝒢_*j*_ is defined as follows:

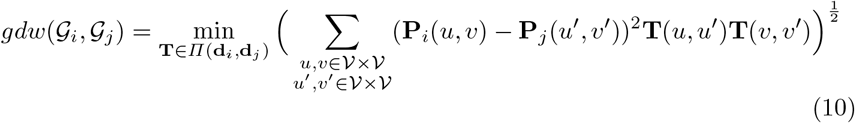

Here, **T** ∈ ℝ^*n*×*n*^ is a transport schedule between the nodes of 𝒢_*i*_ and 𝒢_*j*_. The constraints are specified so that the total transport to/from each node is equal to degree of the node in its respective network, i.e., **d**_*i*_(*u*) = Σ_*v*∈*ν*_**P**_*i*_(*u, v*)*/n* for all *u* ∈ 𝒱, similarly for **d**_*j*_, and *Π*(**d**_*i*_, **d**_*j*_) {**T** ≥ **0** : **T1** = **d**_*i*_, **T**^*T*^ **1** = **d**_*j*_}.

The optimal solution to this problem is a fuzzy matching of the nodes of the two layers such that the nodes with similar topological profiles are mapped to each other. The value of the optimal solution is a measure of the topological dissimilarity between the two networks. Note that there is one-to-one mapping between the nodes of the networks compared in the context of our problem. However, it is still useful to allow mapping of different nodes to each other, since biological networks generally exhibit modular structure [15], thus functionally/biologically similar nodes can compensate for missing edges of each other [13].

For a pair of networks 𝒢_*i*_ and 𝒢_*j*_, *gdw*(𝒢_*i*_, 𝒢_*j*_) can be computed using proximal point algorithm [6]. We compute the weight *w*_*ij*_ of 𝒢_*i*_ and 𝒢_*j*_ based on this value:

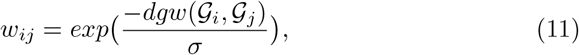

where *σ* is a normalization parameter. This formulation assigns a larger weight to pairs of networks that are topologically similar to each other.

## 3 Results and Discussion

### 3.1 Datasets and Experimental Setup

In our computational experiments, we use datasets that are curated for drug repositioning and drug target prediction, and have been previously used to benchmark machine learning algorithms in the context of these problems [27,26]. These problems are illustrated in the Supplementary Material. The descriptive statics of the datasets are shown in Table 1.

**Table 1:** Descriptive statics of the networks used in computational experiments. Upper table: Bipartite graphs that contain the associations to be predicted. Lower table: Multiplex networks that are used to compute node embeddings, which in turn are used in predictions. Shaded and white rows respectively show networks used in drug-target prediction and drug repositioning.

| Dataset | # Drugs | # Diseases | # Proteins | # Edges |
| --- | --- | --- | --- | --- |
| Drug-Target Network | 1519 | 1229 | – | 6677 |
| Drug-Protein Network | 732 | – | 1915 | 1208 |

Table 1: **Descriptive statics of the networks used in computational experiments.**
| Dataset | # Layers | # Nodes | # Edges |
| --- | --- | --- | --- |
| Large Multiplex Drug Network | 9 | 1519 | [40227–1152921] |
| Small Multiplex Drug Network | 9 | 732 | [66018–267546] |
| Multiplex PPI Network | 6 | 1915 | [11131–1832554] |

#### Drug Repositioning

The task we consider is to predict drug-disease associations based on existing knowledge of drug-disease associations and drug-drug associations. The dataset contains 6677 clinically validated drug-disease associations among 1519 drugs and 1229 diseases [27]. We use a drug multiplex network with 9 layers: (i) clinically reported drug-drug interactions, (ii) drug-target interactions, (iii) drug side-effect associations, (iv) chemical similarity of the drugs, (v) therapeutic similarity the drugs, (vi) sequence similarity of drug targets, and the functional similarity of drug targets based on Gene Ontology (GO) (vii) biological process, (viii) cellular component, (ix) and molecular function [27]. Note that some of these networks are quite dense as they assign a similarity score to all pairs of drugs and do not apply any threshold to sparsify the graph.

#### Drug-Target Interaction Prediction

The task we consider is to predict drug-target interactions based on existing knowledge of drug-target (protein) interactions, drug-drug associations/interactions, and protein-protein interactions. The drug-target interaction network is a bipartite graph consisting of 1208 drug-target pairs among 732 drugs and 1915 proteins/targets [27], obtained from DrugBank, Therapeutic Target Database (TTD), and PharmGKB. The drug multiplex network is a subset of the above network and contains nine layers. The protein multiplex network consists of six layers: (i) STRING PPI, (ii) similarity of the disease-association profiles of proteins, (iii) protein sequence similarities, and functional similarity between proteins quantified based on Gene Ontology (iv) biological process, (v) cellular component, and (vi) molecular function. Similar to the drug networks, some of the protein networks are highly dense.

#### Implementation Details and Competing Algorithms

To measure the Gromov Wassertein Discrepancy (GWD) among layers of multiplex networks, we build our Python code on the implementation provided by [24]. We use this tool with *σ* = 100 to compute *w*_*ij*_’s for all multiplex networks. The weight matrices for each of the three multiplex networks are shown in Supplementary Materials. Since we observe that drug side-effect association and therapeutic similarity networks are considerably dissimilar to other layers, we also consider Pruned Hattusha, a version that discards these two layers in our experiments. For protein-protein interaction networks, all layers exhibit similar topology, thus we retain all layers of the mutiplex PPI network. We implement Hattusha in Matlab and compute 512-dimensional node embeddings by setting *α* = 0.85. Our results on the effect of *α* and the number of embedding dimensions are provided in Supplementary Materials.

We compare Hattusha against two state-of-the-art multiplex network embedding algorithms illustrated in Figure 1, MASHUP [5] and deepDR [27]. We also consider a special case of Hattusha, where we set *w*_*ij*_ = 1 (i.e., all networks are considered equally). This can also be considered an extension of the RWR-M algorithm [23], which is not developed for network embedding, but can be used for network embedding by applying our framework for integrating diffusion states. We refer to this third competing algorithm as RWR-M.

Once drug embeddings are computed, the drug repositioning problem becomes a recommendation problem where drug embeddings are features and drugs are “labels” to be predicted. To accomplish learning and prediction in this setting, we use collective variational autoencoder (cVAE) and its variant fVAE, which are both learning algorithms developed in the context of recommendation systems [4]. In our cross-validation setting (described below), we consider the 6677 drug-disease associations as positively labeled examples and randomly sample 6677 negatively-labeled examples from the 1519×1229−6677 drug-disease pairs that do not have reported association.

The drug-target prediction problem is a supervised link prediction problem, where both drug embeddings and protein (target) embeddings are available. To train and test classifiers for this problem, we use Deep Random Forest [2,27] and Logistic Regression [8] on the concatenated embeddings of drug-protein pairs. In our cross-validation setting (described below), we consider the 1915 drug-target associations as positively labeled examples and randomly sample 1915 negatively-labeled examples from the 1208×1208 − 1915 drug-target pairs that do not have reported association.

In all experiments, we perform 5-fold cross validation and report the mean and standard deviation Area Under ROC Curve (AUC) and Area Under Precision Curve (AUPRC) across five cross-validation runs. The Python and Matlab implementations of our framework are available at https://github.com/mustafaCoskunAgu/Hattusha.

### 3.2 Predictive Accuracy of Multiplex Network Embeddings

The comparison of the predictive accuracy of embeddings provided by Hattusha against three competing algorithms in the context of Drug Repositioning is shown in Figure 2. As seen in the figure, Hattusha (represented by dark blue boxes) delivers best area under ROC curve (AUROC) across the board and outperforms the other three algorithms for all values of *β* we consider. Among the three competing algorithms, MASHUP delivers best performance after Hattusha, followed by RWR-M. This observation suggests that linear and/or probabilistic dimensionality reduction techniques are more effective than neural networks in computing embeddings from diffusion states. The pruned version of Hattusha, which prunes out 2 layers of the drug-drug association network and retains 7 layers, delivers similar AUC as Hattusha, but exhibits significant improvement when area under precision curve (AUPRC) is considered. This suggests that multiplex network layers that have distinct topology as compared to other layers may create confusion for predictive tasks. Finally, although the models that use cVAE consistently perform better than models that use fVAE, the performance difference between embedding algorithms is consistent regardless of the underlying machine learning algorithm.

**Fig 2:**
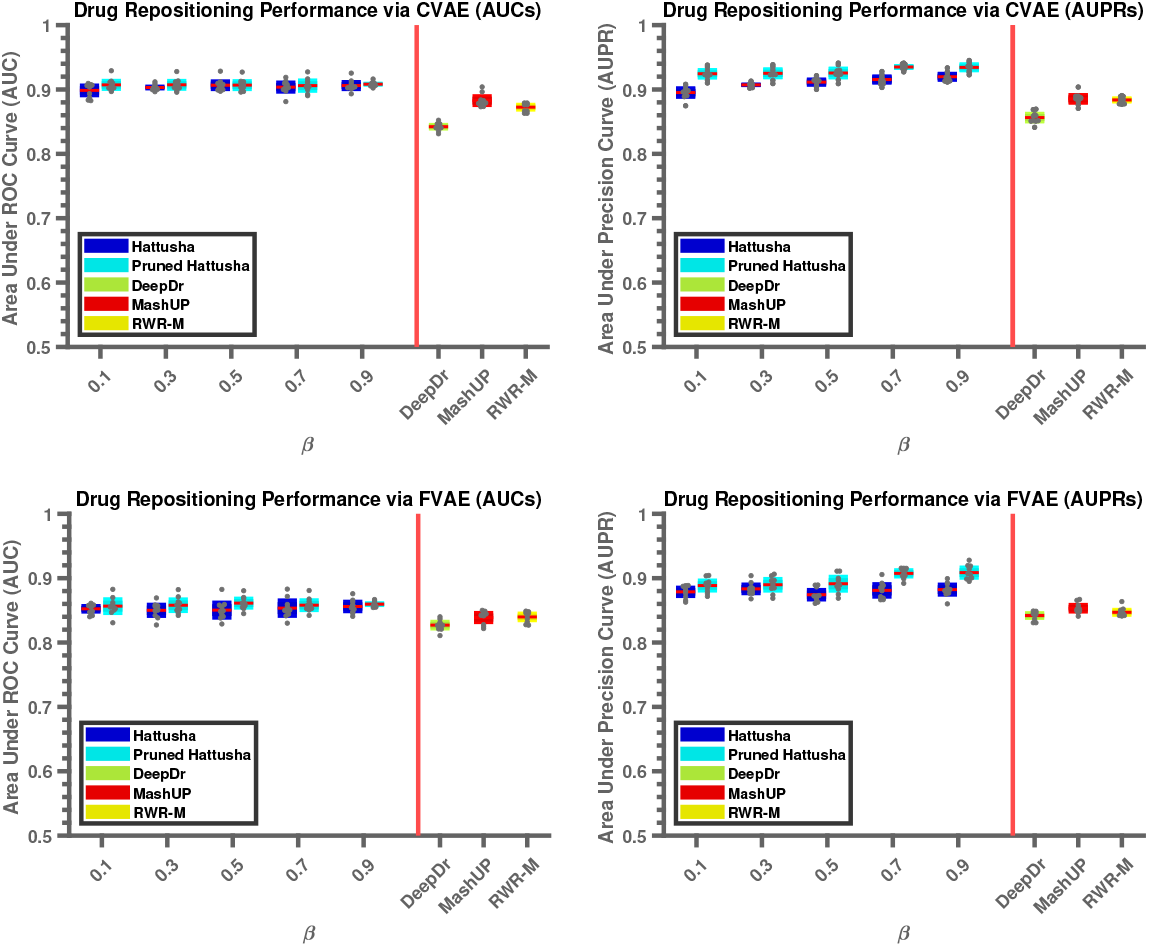
Comparison of predictive accuracy in the context of Drug Repositioning. The mean and standard deviation of the AUROC (left) and AUPRC (right) of models trained using CVAE (top) and FVAE (bottom) are shown for the four embedding algorithms. The performance of Hattusha is shown as a function of *β*, the parameter that acontrols inter-network smoothness of diffusion states. Hattusha refers to the version that uses all layers of the multiplex network, whereas Pruned Hattusha refers to the version that only uses the layers that are most consistent with other layers.

The comparison of the predictive accuracy of embeddings provided by Hattusha against three competing algorithms in the context of Drug-Target Prediction is shown in Figure 3. As seen in the figure, the embeddings computed by Hattusha outperform the embeddings computed by DEEPDR and MASHUP for drug-target prediction as well, regardless of the machine learning algorithm used. However, the performance difference is less pronounced as compared to the drug repositioning problem. Furthermore, RWR-M delivers predictive performance that is very close to that of Hattusha and the improvement provided by pruning out the two dissimilar drug-drug association networks is minimal. These observations further demonstrate the value of accounting for the topological similarity of network layers, as the layers of the multiplex PPI network are topologically similar to each other across the board (Supplementary Figure 1).

**Fig 3:**
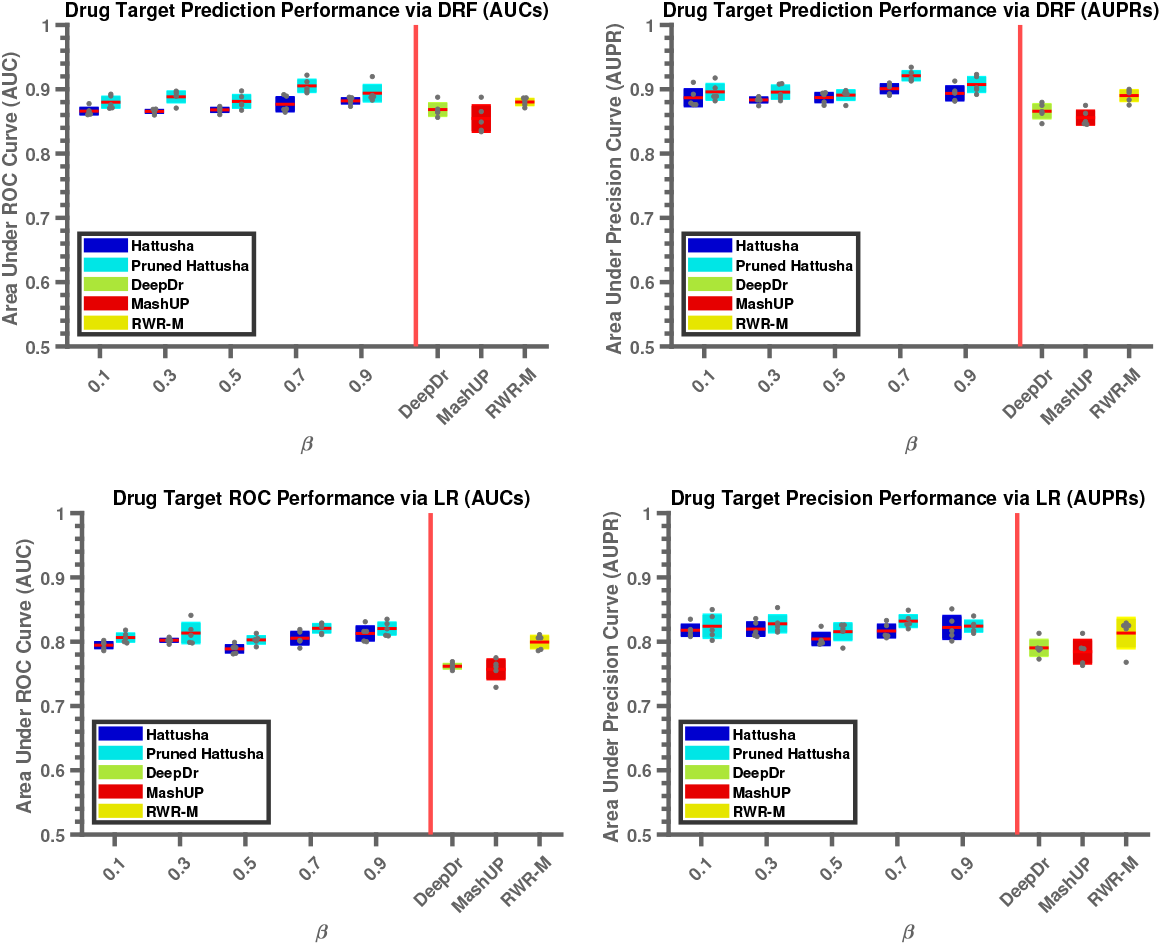
Comparison of predictive accuracy in the context of Drug Target Prediction. The mean and standard deviation of the AUC (left) and AUPR (right) of models trained using Deep Random Forest (top) and Logistic Regression (bottom) are shown for the embeddings provided by the four embedding algorithms. The performance of Hattusha is shown as a function of *β*, the parameter that adjusts the importance of inter-network smoothness of diffusion states. Hattusha refers to the version that uses all layers of the multiplex network, whereas Pruned Hattusha refers to the version that only uses the layers that are most consistent with other layers.

## 4 Conclusion

In this paper, we propose a computational algorithm, Hattusha, for integrating and embedding multiplex biological networks by taking into account the topological similarity of different network layers. Our results show that the topological similarities computed with this measure help improve the predictive performance of the resulting embeddings, while also enabling identification and removal of layers with distinct topological profiles. Future efforts in this direction would include extension of our framework to the integration of heterogeneous networks, as well as other biological applications.

## 1 Supplementary Methods

### 1.1 Efficient Computation of Diffusion States

The optimization problem we formulate in Hattusha can be solved using the following update rule:

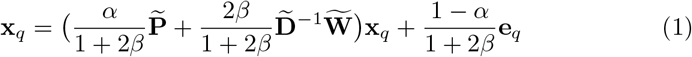

While this update rule lends itself to an iterative solution with standart power iteration, this is computationally expensive as it involves a very large linear system. For this reason, we use Chebyshev acceleration to implement this iterative solution. The idea behind the Chebyshev acceleration technique is to use the summation of all values of the iterate (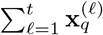, where *t* denotes the current iteration) across all iterations, as opposed to the iterate itself 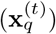. This enables faster convergence, as historical values of the iterate also provide information on its future values (i.e., in the random walk setting, where the random walker has been provides information on where it is headed). Formally, it can be shown that the residual can be bounded by an exponential function of the number of iterations, such that the base of the exponent is strictly smaller than the base associated with the standard power iteration [2].

The iterate for Chebyshev acceleration can be computed using only the last three values of the iterate (namely 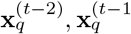, and 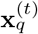) provided that the smallest and largest eigenvalues of the coefficient matrix associated with the system are known and are in the interval [−1, 1] [3]. As stated by the below lemma, this is the case for the system of Equation (1), thus Cheybyshev acceleration can be applied to our problem without requiring extra space. The resulting algorithm is shown in Algorithm 1.

#### Lemma 1.

*The smallest and largest eigenvalues of the coefficient matrix of Equation 1*, 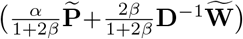, *are respectively* 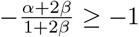 *and* 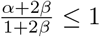.

#### Algorithm 1: Hattusha

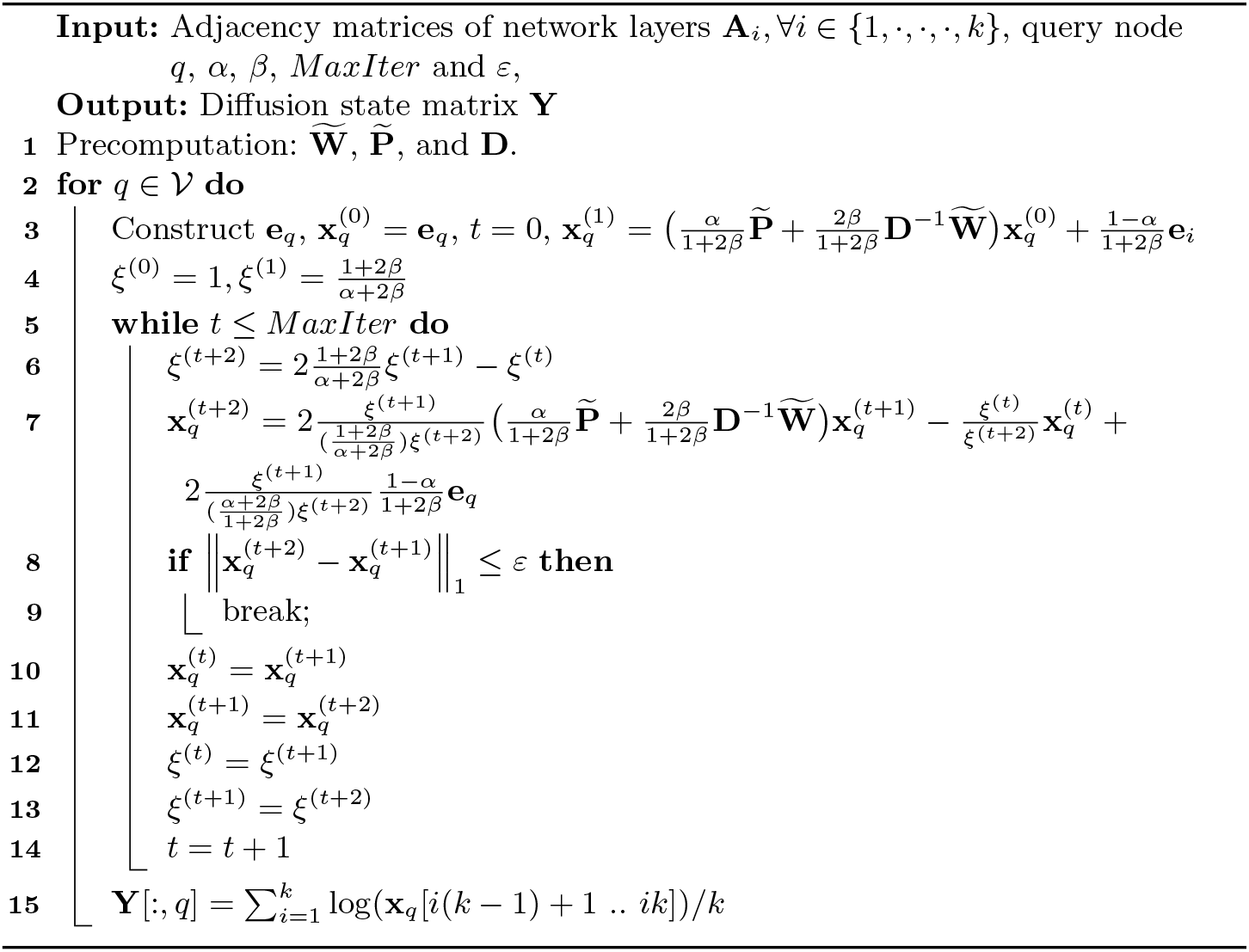

*Proof*. As all networks in each layer of multiplex network are (undirected) symmetric, 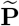 is a positive and symmetric matrix. Also, it is known that degree normalized symmetric positive definite matrix has real eigenvalues in [−1, 1] [4].

Furthermore, from the construction of 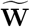 is symmetric positive definite (SPD) as ∀*ij, w*_*ij*_ = *w*_*ji*_ ∈ ℝ^+^ as *exp* : ℝ → ℝ^+^ and 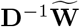 is a stochastic matrix with eigenvalues in [−1, 1] interval.

Let *λ*_1_ and *λ*_*n*_ denotes the largest and smallest eigenvalue of an SPD matrix. Then, from Wesly Theorem [1], we can write, 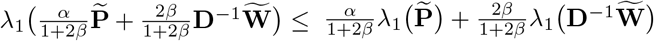

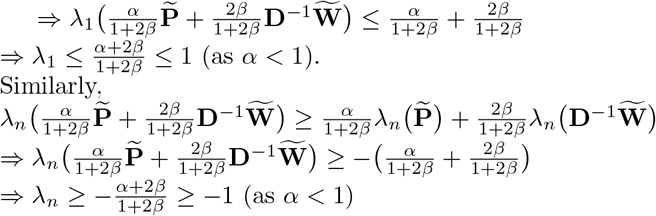

**Supplementary Fig 1:**
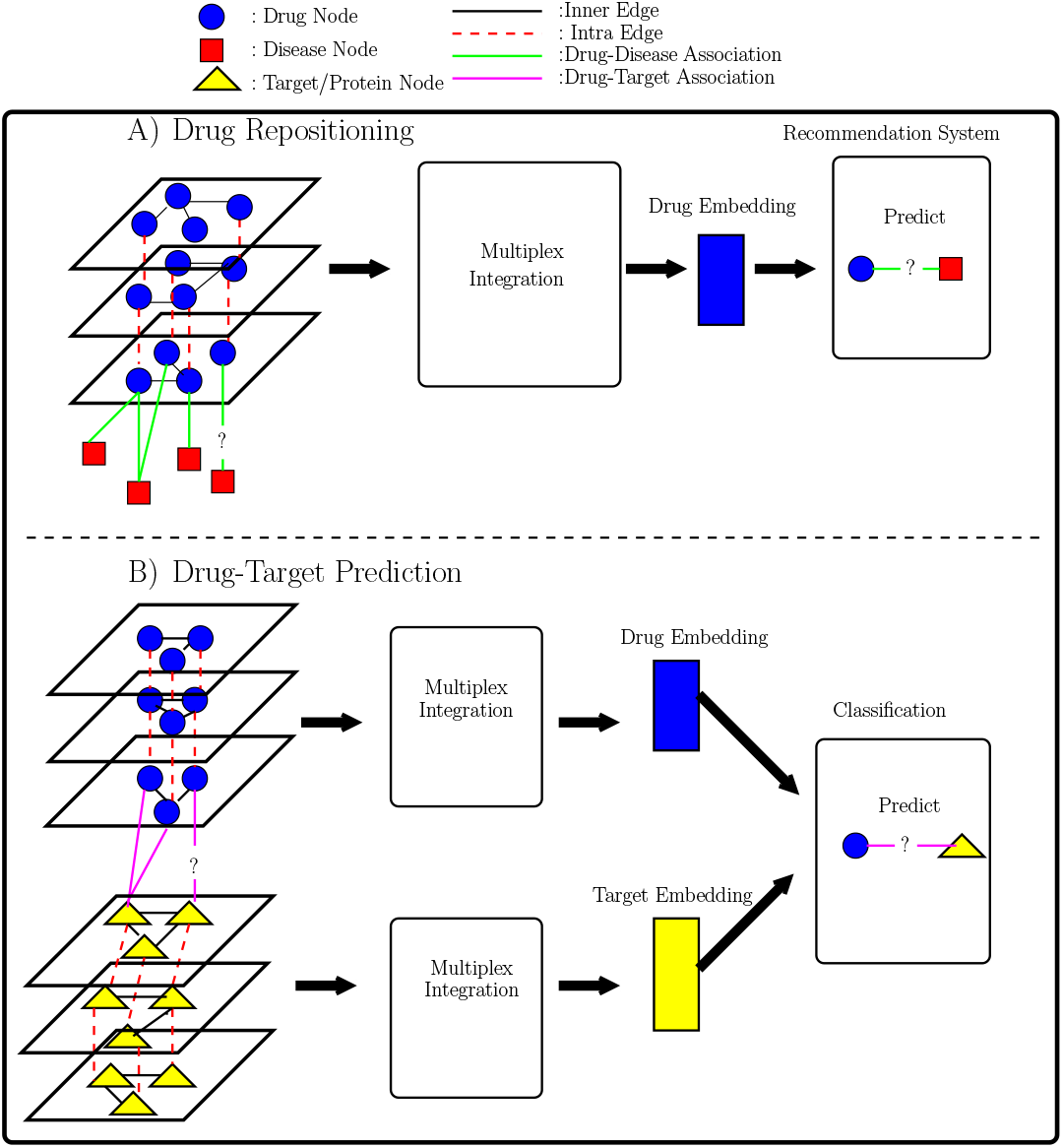
Experimental Setup for the Drug Repositioning and Drug Target Prediction Problems.

## 2 Supplementary Experimental Results

The experimental setup for the two problems we consider are shown in Supplementary Figure 1.

### 2.1 Drug-Drug Networks

The drug-drug association networks are the following:

– DDI: clinically reported drug-drug interactions
– DTI: drug-target interactions
– DSEA: drug side-effect associations
– DChS: chemical similarity of the drugs
– DThS: therapeutic similarity the drugs
– DTSeS: sequence similarity of drug targets,
– DTBPS: functional similarity of drug targets based on Gene Ontology (GO) biological process
– DTCCS: functional similarity of drug targets based on GO cellular component
– DTMFS: functional similarity of drug targets based on GO molecular function

### 2.2 Protein-Protein Networks

The protein-protein interaction/association networks are the following:

– PPI: STRING PPI
– PDA: similarity of the disease-association profiles of proteins
– PSes: protein sequence similarities
– PBPS: functional similarity between proteins quantified based on GO biological process
– PCCS: functional similarity between proteins quantified based on GO cellular component
– PMFS: functional similarity between proteins quantified based on GO molecular function

The weight matrix for the three multiplex networks used in our computational experiments are shown in Supplementary Figure 2.

**Supplementary Fig 2:**
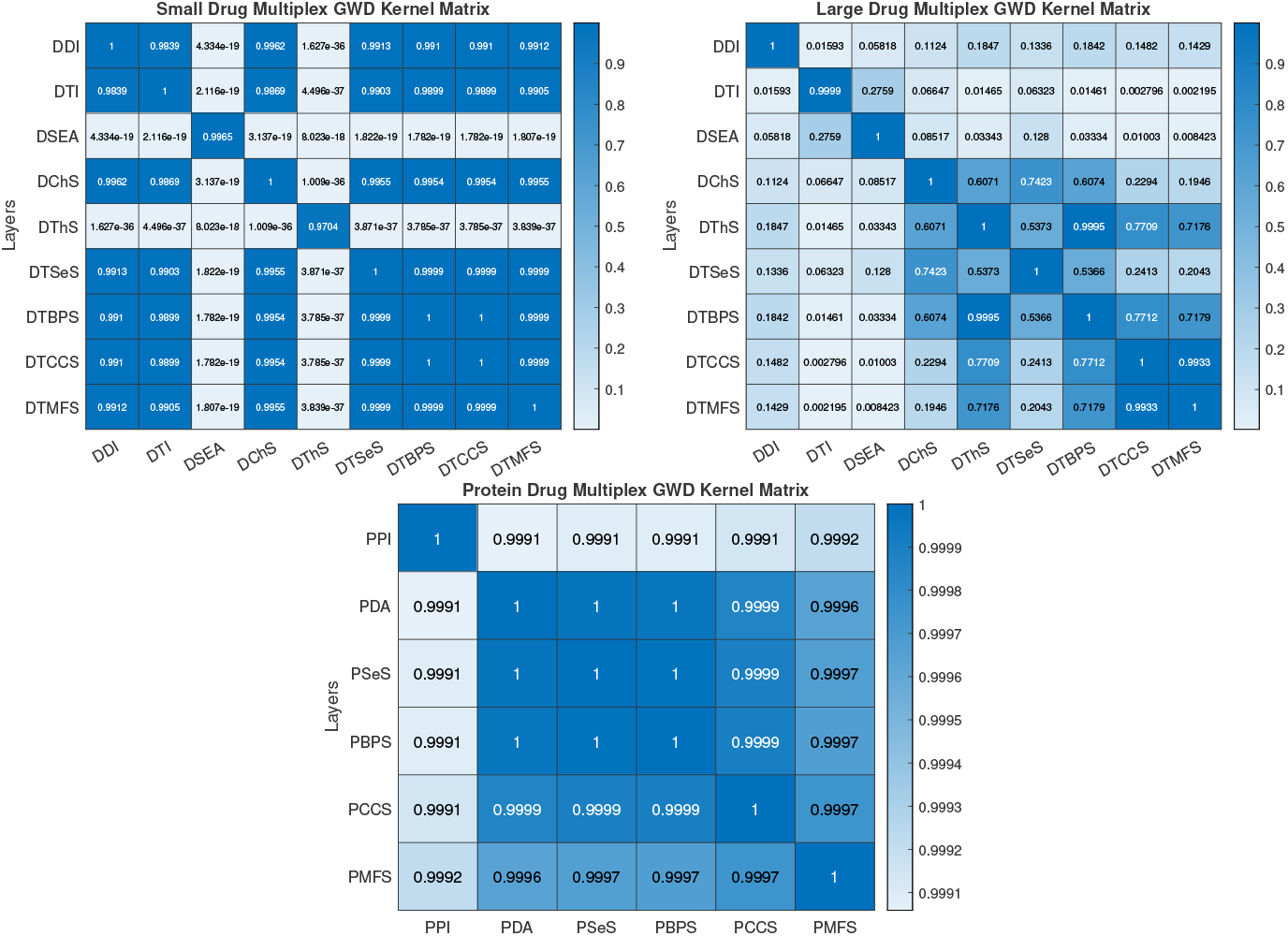
Heatmap of the weights (topological similarity) of network pairs. Each heatmap shows the weight assigned to pairs of multiplex network layers based on Gromov-Wasserteins discrepancy. Darker blue indicates higher topological similarity between two networks.

**Supplementary Fig 3:**
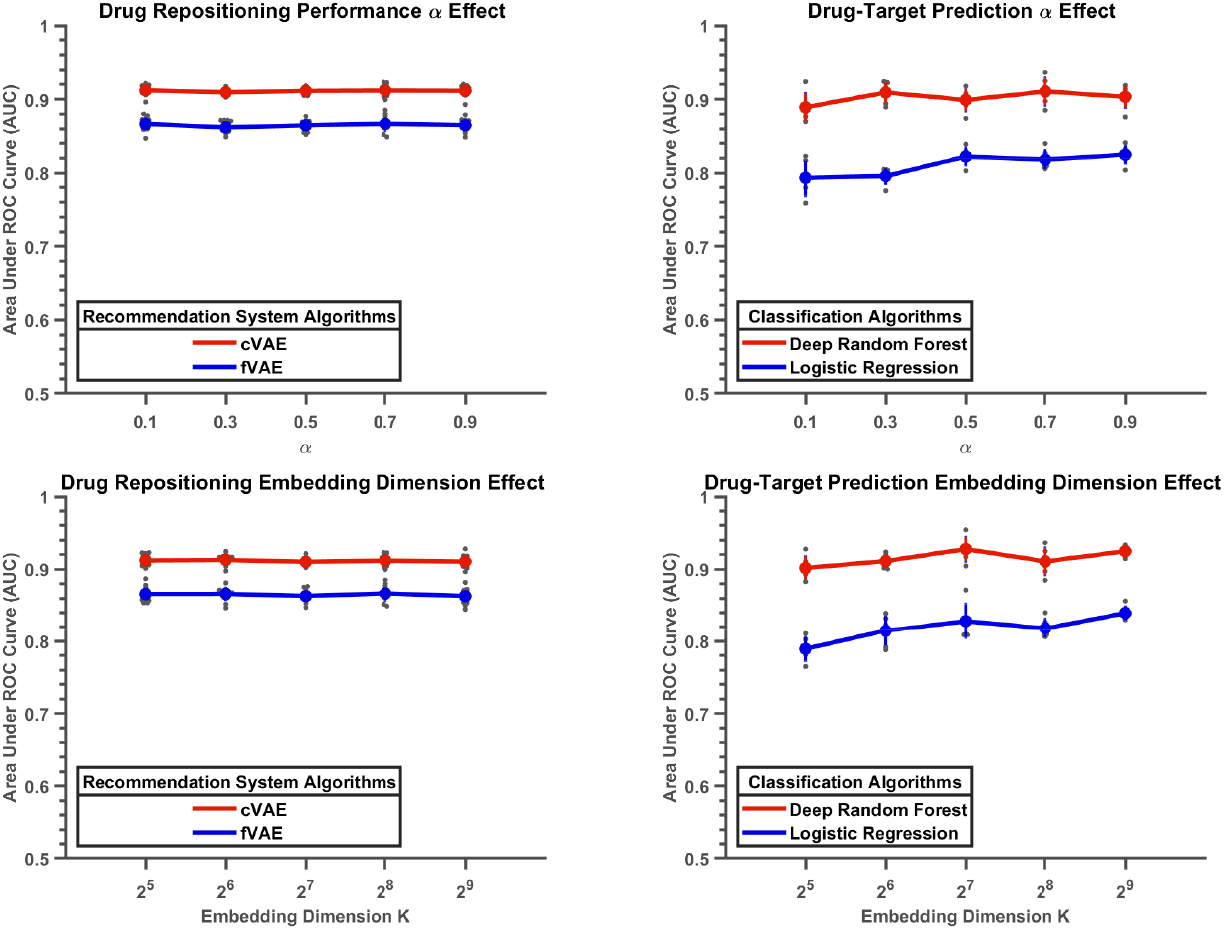
The effect of hyper-parameters on predictive performance. The AUC of models built using embeddings computed by Hattusha in the context of Drug Repositioning (left) and Drug-Target Prediction (right) are shown as a function of the damping factor controlling inner-network smoothness (*α*, top) and number of embedding dimensions (bottom).

### 2.3 Effect of Hyper-parameters

In addition to *β*, which is used to control the importance of inter-network smoothness, there are two additional hyper-parameters that can influence the quality of embeddings: (i) *α*, the damping factor that is is used to control the importance of intra-network smoothness, and (ii) the number of embedding dimensions. We investigate the effect of these parameters by setting *β* = 0.7. In the experiments for *α*, we set the number of embedding dimensions to 512. In the experiments for the number of dimensions, we set *α* = 0.7. The results of this analysis are shown in Figure 3. As seen in the figure, both parameters have little effect on predictive performance for the drug repositioning problem. For the drug-target prediction problem, however, predictive accuracy improves as the number of embedding dimensions, as well as the importance of intra-network smoothness go up, and stabilize/start slowly declining after some point.

### 2.4 Runtime Analysis

Finally, we compare the performance of Chebyshev based efficient multiple network integration, Algorithm 1, with an iterative approach, power method, in terms of running time as a function of *β*. For this purpose, we randomly generate 1000 queries for each multiplex network and report average runtimes of both algorithms in second as seen in the Figure 4. It can be seen in the plots Chebyshev based acceleration renders the computation more efficient than power method, especially for larger *β*, which yields faster embedding for all three multiplex networks that are considered.

**Supplementary Fig 4:**
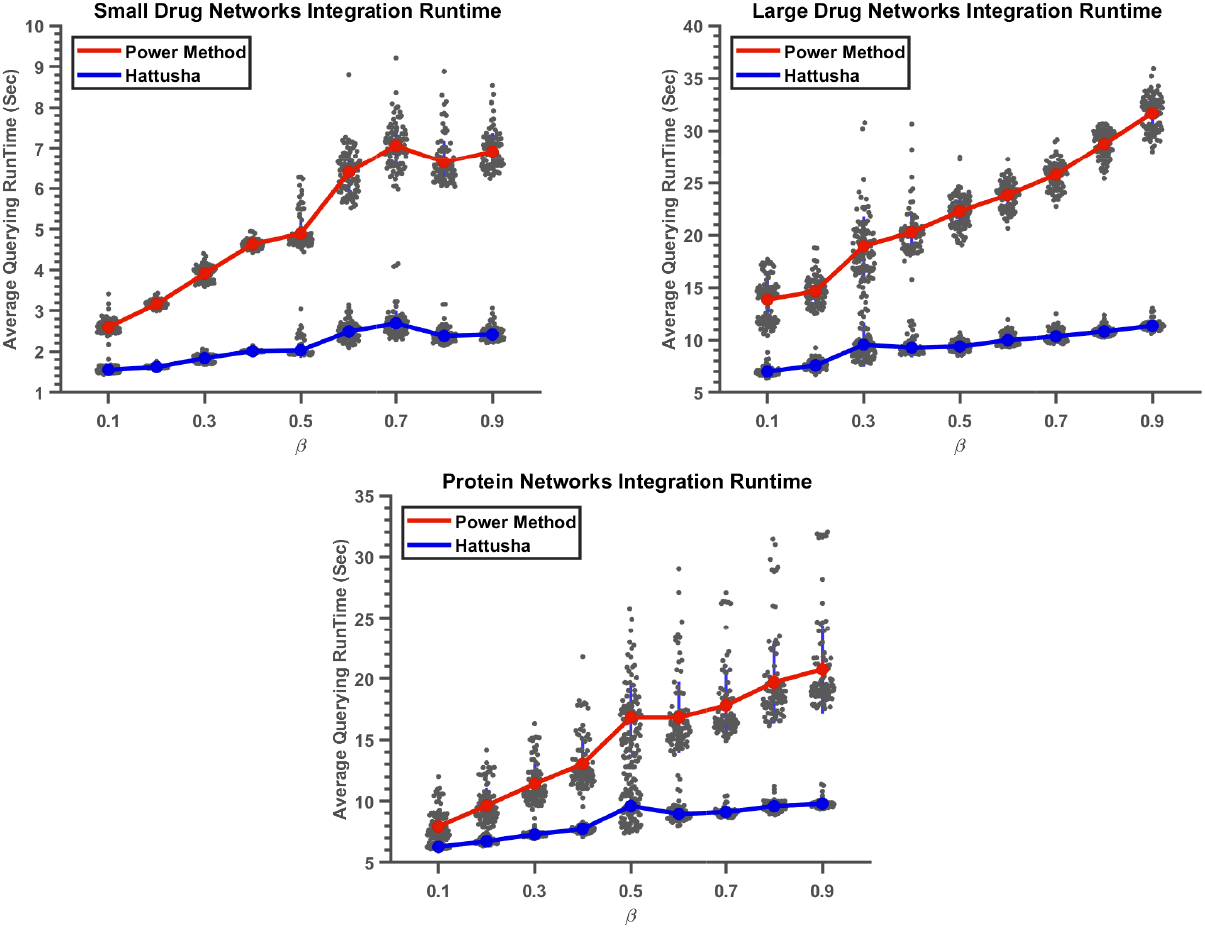
Runtime analysis of multiplex network integration as a function of *β*. We choose 1000 random query nodes, drugs, and report their average query time as a function of *β*.

## Notes

https://github.com/mustafaCoskunAgu/Hattusha

### Competing Interest Statement

The authors have declared no competing interest.

## References

1. Albert, I., Albert, R.: Conserved network motifs allow protein–protein interaction prediction. Bioinformatics 20(18), 3346–3352 (2004)

2. Breiman, L.: Random forests. Machine learning 45(1), 5–32 (2001)

3. Camacho, D.M., Collins, K.M., Powers, R.K., Costello, J.C., Collins, J.J.: Next-generation machine learning for biological networks. Cell 173(7), 1581–1592 (2018)

4. Chen, Y., de Rijke, M.: A collective variational autoencoder for top-n recommendation with side information. In: Proceedings of the 3rd Workshop on Deep Learning for Recommender Systems. pp. 3–9 (2018)

5. Cho, H., Berger, B., Peng, J.: Compact integration of multi-network topology for functional analysis of genes. Cell systems 3(6), 540–548 (2016)

6. Chowdhury, S., Mémoli, F.: The gromov–wasserstein distance between networks and stable network invariants. Information and Inference: A Journal of the IMA 8(4), 757–787 (2019)

7. Coşkun, M., Grama, A., Koyutürk, M.: Efficient processing of network proximity queries via chebyshev acceleration. In: Proceedings of the 22nd ACM SIGKDD International conference on knowledge discovery and data mining. pp. 1515–1524 (2016)

8. Coşkun, M., Koyutürk, M.: Node similarity based graph convolution for link prediction in biological networks (2021)

9. Cowen, L., Ideker, T., Raphael, B.J., Sharan, R.: Network propagation: a universal amplifier of genetic associations. Nature Reviews Genetics 18(9), 551 (2017)

10. De Las Rivas, J., Fontanillo, C.: Protein–protein interactions essentials: key concepts to building and analyzing interactome networks. PLoS computational biology 6(6), e1000807 (2010)

11. Gomez, S., Diaz-Guilera, A., Gomez-Gardenes, J., Perez-Vicente, C.J., Moreno, Y., Arenas, A.: Diffusion dynamics on multiplex networks. Physical review letters 110(2), 028701 (2013)

12. Li, M., Koyutürk, M.: Consensus embeddings for networks with multiple versions. In: International Conference on Complex Networks and Their Applications. pp. 39–52. Springer (2020)

13. Liao, C.S., Lu, K., Baym, M., Singh, R., Berger, B.: Isorankn: spectral methods for global alignment of multiple protein networks. Bioinformatics 25(12), i253–i258 (2009)

14. Lonsdale, J., Thomas, J., Salvatore, M., Phillips, R., Lo, E., Shad, S., Hasz, R., Walters, G., Garcia, F., Young, N., et al.: The genotype-tissue expression (gtex) project. Nature genetics 45(6), 580–585 (2013)

15. Mitra, K., Carvunis, A.R., Ramesh, S.K., Ideker, T.: Integrative approaches for finding modular structure in biological networks. Nature Reviews Genetics 14(10), 719–732 (2013)

16. Nelson, W., Zitnik, M., Wang, B., Leskovec, J., Goldenberg, A., Sharan, R.: To embed or not: network embedding as a paradigm in computational biology. Frontiers in genetics 10, 381 (2019)

17. Ni, J., Tong, H., Fan, W., Zhang, X.: Inside the atoms: ranking on a network of networks. In: Proceedings of the 20th ACM SIGKDD international conference on Knowledge discovery and data mining. pp. 1356–1365 (2014)

18. Park, C., Kim, D., Han, J., Yu, H.: Unsupervised attributed multiplex network embedding. In: Proceedings of the AAAI Conference on Artificial Intelligence. vol. 34, pp. 5371–5378 (2020)

19. Peyré, G., Cuturi, M., Solomon, J.: Gromov-wasserstein averaging of kernel and distance matrices. In: International Conference on Machine Learning. pp. 2664–2672. PMLR (2016)

20. Roweis, S.T., Saul, L.K.: Nonlinear dimensionality reduction by locally linear embedding. science 290(5500), 2323–2326 (2000)

21. Ruiz, C., Zitnik, M., Leskovec, J.: Identification of disease treatment mechanisms through the multiscale interactome. Nature communications 12(1), 1–15 (2021)

22. Su, C., Tong, J., Zhu, Y., Cui, P., Wang, F.: Network embedding in biomedical data science. Briefings in bioinformatics 21(1), 182–197 (2020)

23. Valdeolivas, A., Tichit, L., Navarro, C., Perrin, S., Odelin, G., Levy, N., Cau, P., Remy, E., Baudot, A.: Random walk with restart on multiplex and heterogeneous biological networks. Bioinformatics 35(3), 497–505 (2019)

24. Vayer, T., Chapel, L., Flamary, R., Tavenard, R., Courty, N.: Fused gromov-wasserstein distance for structured objects. Algorithms 13(9), 212 (2020)

25. Yue, X., Wang, Z., Huang, J., Parthasarathy, S., Moosavinasab, S., Huang, Y., Lin, S.M., Zhang, W., Zhang, P., Sun, H.: Graph embedding on biomedical networks: methods, applications and evaluations. Bioinformatics 36(4), 1241–1251 (2020)

26. Zeng, X., Zhu, S., Hou, Y., Zhang, P., Li, L., Li, J., Huang, L.F., Lewis, S.J., Nussi-nov, R., Cheng, F.: Network-based prediction of drug–target interactions using an arbitrary-order proximity embedded deep forest. Bioinformatics 36(9), 2805–2812 (2020)

27. Zeng, X., Zhu, S., Liu, X., Zhou, Y., Nussinov, R., Cheng, F.: deepdr: a network-based deep learning approach to in silico drug repositioning. Bioinformatics 35(24), 5191–5198 (2019)

## References

1. Bhatia, R.: Linear algebra to quantum cohomology: the story of alfred horn’s inequalities. The American Mathematical Monthly 108(4), 289–318 (2001)

2. Coskun, M., Grama, A., Koyuturk, M.: Efficient processing of network proximity queries via chebyshev acceleration. In: Proceedings of the 22nd ACM SIGKDD International conference on knowledge discovery and data mining. pp. 1515–1524 (2016)

3. Saad, Y.: Chebyshev acceleration techniques for solving nonsymmetric eigenvalue problems. Mathematics of Computation 42(166), 567–588 (1984)

4. Saad, Y.: Iterative methods for sparse linear systems. SIAM (2003)

